# Single molecule, full-length transcript sequencing provides insight into the extreme metabolism of ruby-throated hummingbird *Archilochus colubris*

**DOI:** 10.1101/117218

**Authors:** Rachael E. Workman, Alexander M. Myrka, Elizabeth Tseng, G. William Wong, Kenneth C. Welch, Winston Timp

**Author notes:** Co-first author (Rachael E. Workman, Alexander M. Myrka).

## Abstract

Hummingbirds can support their high metabolic rates exclusively by oxidizing ingested sugars, which is unsurprising given their sugar-rich nectar diet and use of energetically expensive hovering flight. However, they cannot rely on dietary sugars as a fuel during fasting periods, such as during the night, at first light, or when undertaking long-distance migratory flights, and must instead rely exclusively on onboard lipids. This metabolic flexibility is remarkable both in that the birds can switch between exclusive use of each fuel type within minutes and in that *de novo* lipogenesis from dietary sugar precursors is the principle way in which fat stores are built, sometimes at exceptionally high rates, such as during the few days prior to a migratory flight. The hummingbird hepatopancreas is the principle location of *de novo* lipogenesis and likely plays a key role in fuel selection, fuel switching, and glucose homeostasis. Yet understanding how this tissue, and the whole organism, achieves and moderates high rates of energy turnover is hampered by a fundamental lack of information regarding how genes coding for relevant enzymes differ in their sequence, expression, and regulation in these unique animals. To address this knowledge gap, we generated a *de novo* transcriptome of the hummingbird liver using PacBio full-length cDNA sequencing (Iso-Seq), yielding a total of 8.6Gb of sequencing data, or 2.6M reads from 4 different size fractions. We analyzed data using the SMRTAnalysis v3.1 Iso-Seq pipeline, including classification of reads and clustering of isoforms (ICE) followed by error-correction (Arrow). With COGENT, we clustered different isoforms into gene families to generate *de novo* gene contigs. We performed orthology analysis to identify closely related sequences between our transcriptome and other avian and human gene sets. We also aligned our transcriptome against the *Calypte anna* genome where possible. Finally, we closely examined homology of critical lipid metabolic genes between our transcriptome data and avian and human genomes. We confirmed high levels of sequence divergence within hummingbird lipogenic enzymes, suggesting a high probability of adaptive divergent function in the hepatic lipogenic pathways. Our results have leveraged cutting-edge technology and a novel bioinformatics pipeline to provide a compelling first direct look at the transcriptome of this incredible organism.

## Introduction

Hummingbirds are the only avian group to engage in sustained hovering flight as a means for accessing floral nectar, their primary caloric energy source. While hovering, small hummingbirds, such as the ruby-throated hummingbird (*Archilochus colubris*), achieve some of the highest mass-specific metabolic rates observed among vertebrates (Suarez 1992; Chai and Dudley 1996). Given their specialized, sugar-rich diet, it is not that surprising that hummingbirds are able to fuel this intense form of exercise exclusively by oxidizing carbohydrates (Suarez et al. 1990; Chen and Welch 2014). This energetic feat is also remarkable in that the source of sugar oxidized by flight muscles during hovering is the same sugar ingested in nectar meals only minutes prior (Chen and Welch 2014; Welch et al. 2007). In addition, hummingbirds seem equally adept at relying on either glucose or fructose (the two monosaccharides comprising their nectar (Baker 1975) as a metabolic fuel for flight (Chen and Welch 2014). In doing so, they achieve rates of sugar flux through their bodies that are up to 55 × greater than non-flying mammals (Welch and Chen 2014).

Hummingbird flight is not always a solely carbohydrate-fueled endeavor. Lipids are a more energy dense form of fuel storage, and fasted hummingbirds are as capable of fueling hovering flight via the oxidation of onboard lipid stores as they are dietary sugars (Welch et al. 2007). Lipids are likely the sole or predominant fuel used during overnight periods (Powers et al. 2003). Just as flux of sugar through the hummingbird is extremely rapid, the building of lipid stores from dietary sugar is also rapid when needed. For example, ruby-throated hummingbirds can routinely increase their mass by 15% or more between midday and dusk on a given day (Hou et al. 2015). The ruby-throated hummingbird (*A. colubris*) completes an arduous annual migratory journey from breeding grounds as far north as Quebec in Canada to wintering grounds in Central America (Weidensaul et al. 2013). Hummingbirds are constrained to fueling long distance migratory flights using onboard lipids. In preparing for such flights, hummingbirds rapidly build fat stores prior to departure or at migratory stopover points, increasing their mass by 25-40% in as few as four days (Carpenter et al. 1993; Hou et al. 2015; Hou and Welch 2016).

The ability to switch so completely and quickly between fuel types means these animals possess remarkably exquisite control over rates of substrate metabolism and biosynthesis in the liver, the principal site of lipogenesis in birds (Hermier 1997). While hummingbird liver does indeed exhibit remarkably high activities of lipogenic and other metabolic enzymes (Suarez et al. 1988), the mechanisms underlying high catalytic rates (high catalytic efficiency and/or high levels of enzyme expression) and control over flux (the role of hierarchical versus metabolic control), sensu (ter Kuile and Westerhoff 2001), remain unclear.

Despite long-standing recognition of, and interest in, their extreme metabolism, the lack of knowledge about gene and protein sequences in hummingbirds has limited more detailed and mechanistic analyses. Amplification of hummingbird genetic sequences for sequencing and/or cloning is hampered by the lack of sequence information from closely related groups, making well-targeted primer design difficult. Only two genes have thus far been cloned from any hummingbird: an uncoupling protein (UCP) homolog and insulin (Vianna et al. 2001; Fan et al. 1993). These two studies offer limited insight into what adaptations in hepatopancreatic molecular physiology underlie extreme energy turnover or its regulation. The UCP homolog was cloned from pectoralis (flight muscle) and its functional significance *in vivo* is unclear. The amino acid sequence of hummingbird insulin was found to be largely identical to that from chicken; however, birds are insulin insensitive and lack the insulin-regulated glucose transporter (GLUT) protein GLUT4, making the role of this hormone in the regulation of energy homeostasis in hummingbirds unknown (Welch et al. 2013; Braun and Sweazea 2008; Polakof et al. 2011).

Recently completed sequencing of the Anna's hummingbird (*Calypte anna*) genome provides a powerful new tool in the arsenal of biologists seeking to understand variation in metabolic physiology in hummingbirds and other groups (Jarvis et al. 2015). Despite their extreme catabolic and anabolic capabilities, hummingbirds have the smallest genome among birds (Gregory et al. 2009) and, in general, have among the smallest vertebrate genomes (Hughes and Hughes 1995). Thus, it seems likely that understanding of transcriptional variation, overlaid on top of genetic variation, is crucial to understanding what makes these organisms such elite metabolic performers.

To this end, we produced the first high-coverage transcriptome of any single avian tissue, the liver of the ruby-throated hummingbird, *Archilochus colubris.* Because many of the proteins involved in cellular metabolism are quite large, we collaborated with Pacific Biosciences to generate long-read sequences as these would enhance our ability to identify full coding sequences and multiple encoded isoforms. The primary advantage to the Pacbio Iso-seq methodology is the capability for full-length transcript sequencing, rendering complete mRNA sequences without the need for assembly. This has been demonstrated in previous studies to drastically increase detection of alternative splicing events (Abdel-Ghany et al. 2016). Additionally, full-length sequences greatly enhance the likelihood of detecting novel or rare splice variants, which is crucial for fully characterizing the transcriptomes of lesser studied, non-model organisms such as the hummingbird.

## Results

### First high coverage single-tissue avian transcriptome quality control and validation

The transcriptome described in this manuscript represents the first long read, single tissue avian transcriptome completed at high coverage. Four size fractions (1-2kb, 2-3kb, 3-6kb, 5-10kb) of sample were sequenced on 40 SMRT Cells, producing 440.75 Gb of raw data, 3.4Gb of full-length non-chimeric reads after circular consensus sequence (CCS) generation and filtering for full-length read classification. Of the four size-selected bins, our average CCS length was 1533, 2464, 3650, and 5444 bp, respectively (Figure 1B).

**Figure 1.**
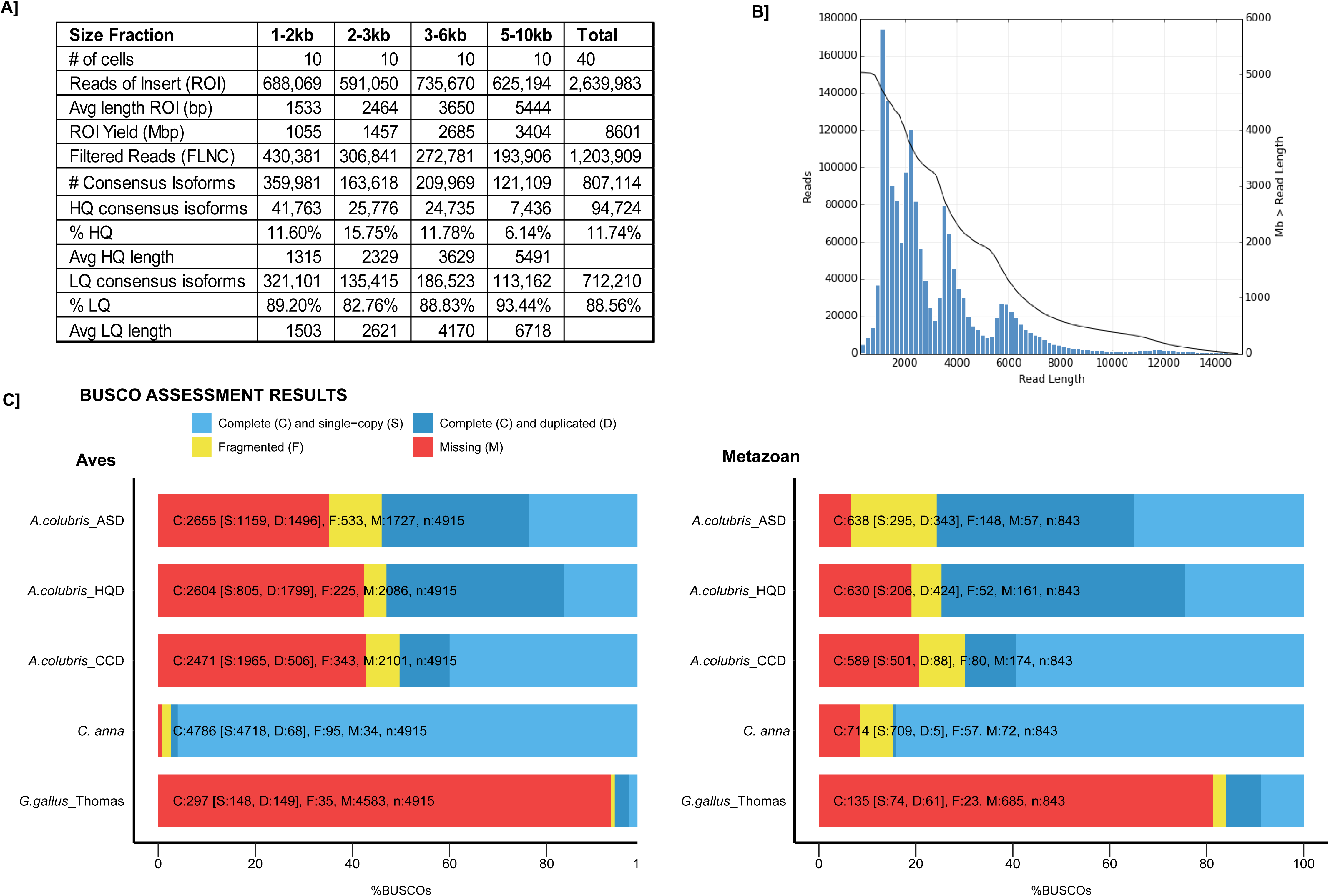
Analysis pipeline. **A** Raw sequence reads from a Pacbio RSII sequencer (bax.h5, bas.h5) were sorted into full and non-full length reads using a classification algorithm that identified full length reads with forward and reverse primers, as well as a poly-A tail. Iterative clustering for isoforms (ICE) was performed on full length reads, and non-full length reads were recruited to perform ARROW polished on the consensus isoforms. Polished sorted reads into high and low-quality bins, and either high quality data (HQD), all sequence data (ASD) or both sets of data, were carried on to further applications (**B**).

To analyse these data, we employed the Pacbio sa3_ds_isoseq pipeline (SMRTAnalysis 3.1.0, (Minoche et al. 2015; Gordon et al. 2015), code here: https://github.com/PacificBiosciences/SMRT-Analysis) using the DNANexus cloud computing platform. A summary of analyses performed using both the SMRTAnalysis pipeline and downstream are displayed in Figure 2A-B. Using this pipeline, we classified 3.4Gb of non-chimeric CCS reads into 1,236,437 full-length (48%) and 1,665,929 non-full length reads where reads were determined to be full-length if both the 5’ and 3’ cDNA primers as well as the polyA tail signal were detected. The Iso-Seq pipeline then performed isoform-level clustering (ICE) followed by final polishing using the Arrow algorithm to output high-quality (predicted accuracy >= 99%), full-length, isoform consensus sequences. The Iso-Seq pipeline produced 238Mb (807,104 reads) of high quality consensus isoforms (HQD, 94,724 reads), and 2Gb (712,210 reads) of low quality consensus isoforms (summary statistics Figure 1A).

**Figure 2.**
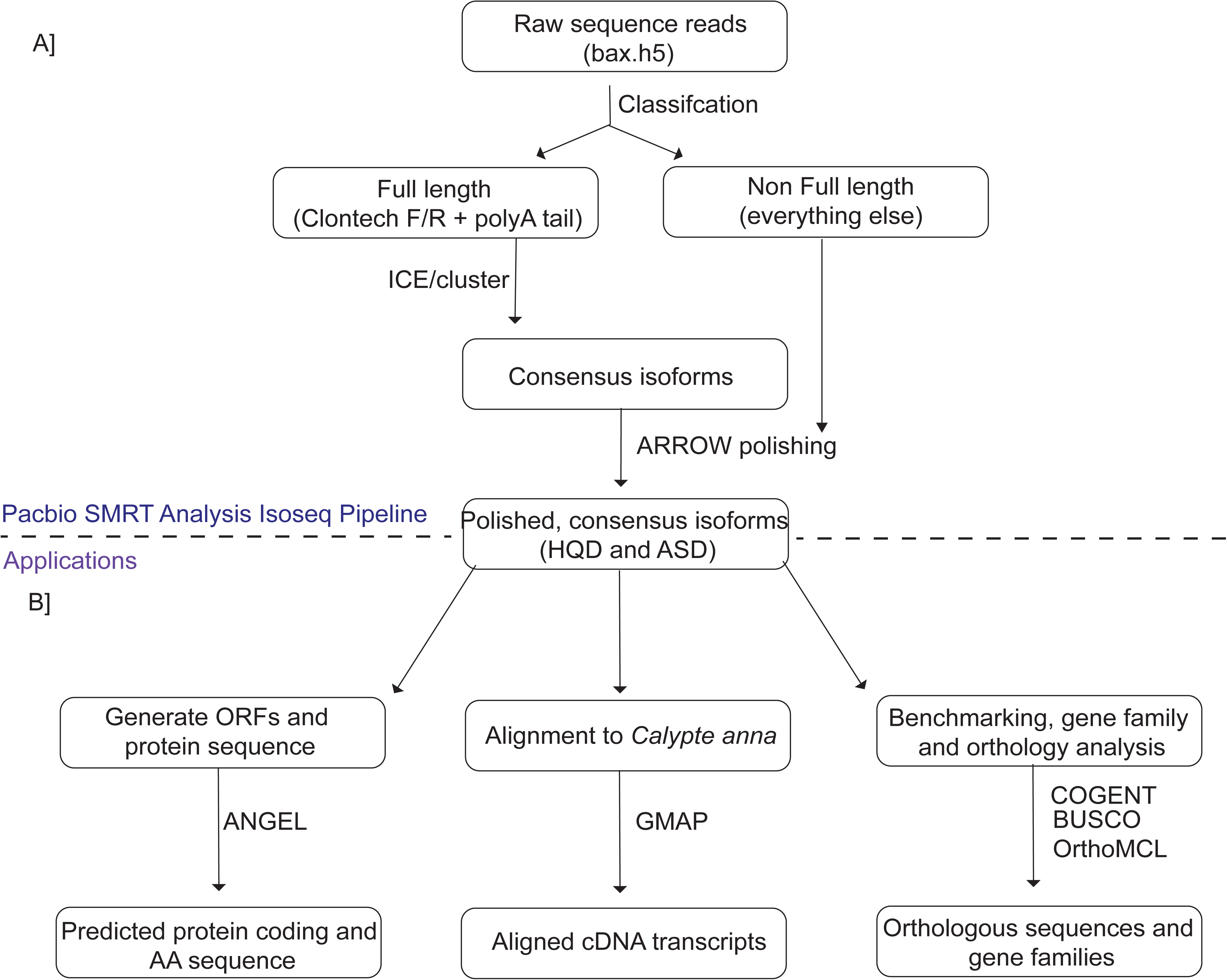
Transcriptome dataset quality control. Average read lengths and isoform counts for 4 sequenced size fractions given in **A**, and read length for all sequence data (ASD, HQ + LQ) plotted in **B**, with black line representing Mb data greater than read length. For example, at 2000bp, 5000Mb of sequence data was larger than 2000bp. **C.** BUSCO transcriptome assessment results for *Archilochus colubris* (ruby-throated hummingbird, all sequence data ASD, high quality sequence data HQD), Cogent-collapsed data (CCD), *Calypte anna* (Anna's hummingbird), *Gallus gallus* (chicken) Thomas (single-tissue transcriptome).

To determine completeness of our transcriptome assembly relative to established transcriptomes, as well as relative retention of transcripts between multiple data processing steps, we used benchmarking software BUSCO (Benchmarking Universal Single Copy Orthologs). BUSCO searches for a list of conserved orthologous genes assumed to be present in all completed transcriptome assemblies for members of the given clade. We utilized the Metazoan and Aves lineage datasets (Simão et al. 2015) to determine assembly completeness for not only our ASD (all-sequence data) dataset, but also the HQD (High quality dataset) and COGENT-collapsed dataset (CCD, described below) in order to ensure that sequence diversity was not lost when filtering. We then compared this result with the chicken transcriptome (*Gallus gallus*) (ftp://ftp.ncbi.nih.gov/genomes/Gallus_gallus), the predicted transcriptome from the recently published Anna's hummingbird (*Calypte anna*) genome (http://gigadb.org/dataset/101004), and *Gallus gallus* from a single-tissue Pacbio Iso-Seq dataset (http://journals.plos.org/plosone/article?id=10.1371/journal.pone.0094650). Busco v2.0 BETA was used (downloaded 9/1/2016, https://gitlab.com/ezlab/busco), and results are summarized in Figure 2C and Supplemental Table 1. Notably, the NCBI *Gallus gallus* transcriptome is nearly complete for both Aves and Metazoan sets, with the predicted transcriptome from *Calypte anna* not far behind. In contrast, our transcriptome contains only about half of the Aves set and slightly more of the metazoan set; many genes are not substantially expressed in a single tissue. Notably our ortholog detection is dramatically improved over the other single-tissue Pacbio data available, (Thomas et al. 2014), which only captured 6% of the Aves set.

To estimate the completeness of our liver transcriptome sequencing, we took subsets of the circular consensus reads dataset and BLASTed against the predicted *Calypte anna* gene set. We found that the number of unique genes detected began to saturate when reaching a 90% subset of our data, suggesting that we had near complete transcriptome sequencing (Supplemental Figure 1). Lower expressed genes may not be detected, but that vast majority of liver expressed genes are likely represented in our data.

### Agreement with established Anna's hummingbird genome reveals general clade conservation

We aligned transcripts to the *Calypte anna* (Anna's hummingbird) genome using both all consensus isoforms (ASD) and high quality isoforms only (HQD) (Korlach et al. 2017), as well as reads collapsed by COGENT (CCD, methods detailed below). As *Calypte anna* is a close relative within the same Bee clade of hummingbirds (McGuire et al. 2007), we expected alignment to perform well. We found an average alignment identity of 94.8%, with 87% transcripts uniquely mapping to the reference. Of the uniquely mapped, 73% covered >90% of the query sequence (alignment length and statistics, Supplemental Figure 2A, 2B), demonstrating high fidelity of aligned reads to reference. When ASD reads were parsed by number of reads of insert supporting each consensus cluster, it was found that generally, alignment identity was high regardless of number of supporting reads. A clear increase in mean alignment identity was found when two or more supporting reads were collapsed (Supplemental Figure 3).

When GMAP was performed using only high quality isoforms (filtered for 2+ full-length supporting reads), alignment percentage was 95.7%, with 93.4% of transcripts mapping uniquely to the reference. The average mapped read length was 2411bp (HQD, 2617bp ASD), while the average predicted CDS length for *Calypte anna* was 1386bp. This being said, reads mapped with GMAP contain UTRs. When we predict just the CDS sequences for *A. colubris* using ANGEL (https://github.com/PacificBiosciences/ANGEL), the mean length was 981bp. When we BLASTed the unaligned reads to whole NCBI database, they largely mapped back to *Calypte anna* (53%). This result suggests that our mapping parameters were too stringent to map these reads, error rate prevented alignment, unaligned regions are divergent enough between both hummingbirds to preclude alignment, or some combination of the above.

### Putative gene family prediction and reduction of transcript redundancy reduces data load while maintaining transcript diversity

To assign transcripts to putative gene families, as well as cluster and eliminate redundant transcripts to produce a unique set of gene isoforms, we utilized the newly developed COGENT (https://github.com/Magdoll/Cogent, Liz Tseng, pre-print pending) pipeline. COGENT is specifically designed for transcriptome assembly in the absence of a reference genome, allowing for isoforms of the same gene to be distinctly identified from different gene families, which are defined as having more than two (possibly redundant) transcript copies. Of the 94,724 HQ consensus isoforms, 91,733 were grouped into 6,725 gene families (Figure 3A). The remaining 2,991 sequences were classified as putative single-isoform genes, or “orphans”. Reconstructed contigs were then applied in place of a reference (or de novo clustering) to reduce redundant transcripts in the original HQD dataset. From this approach, we were able to reduce our HQ dataset to 14,628 distinct transcript isoforms and 2990 orphan isoforms, for a total of 17,618 isoform sequences (18% of the original). Data is further summarized in Supplemental Table 2. An average of 1.53 isoforms was found per gene family (Figure 3B), with 2624, or 27.4% of the gene families having more than one isoform, including “orphans”. While other studies have found more isoforms per locus, for example 6.56 in (Wang et al. 2016), that study multiplexed six plant tissues, whereas a lower complexity is to be expected with single tissue analysis. This dataset (COGENT collapsed data, or CCD) was also mapped onto the *Calypte anna* genome assembly (http://gigadb.org/dataset/101004), to demonstrate the effectiveness of this method in reducing transcript redundancy and classifying isoforms (Figure 3C).

**Figure 3.**
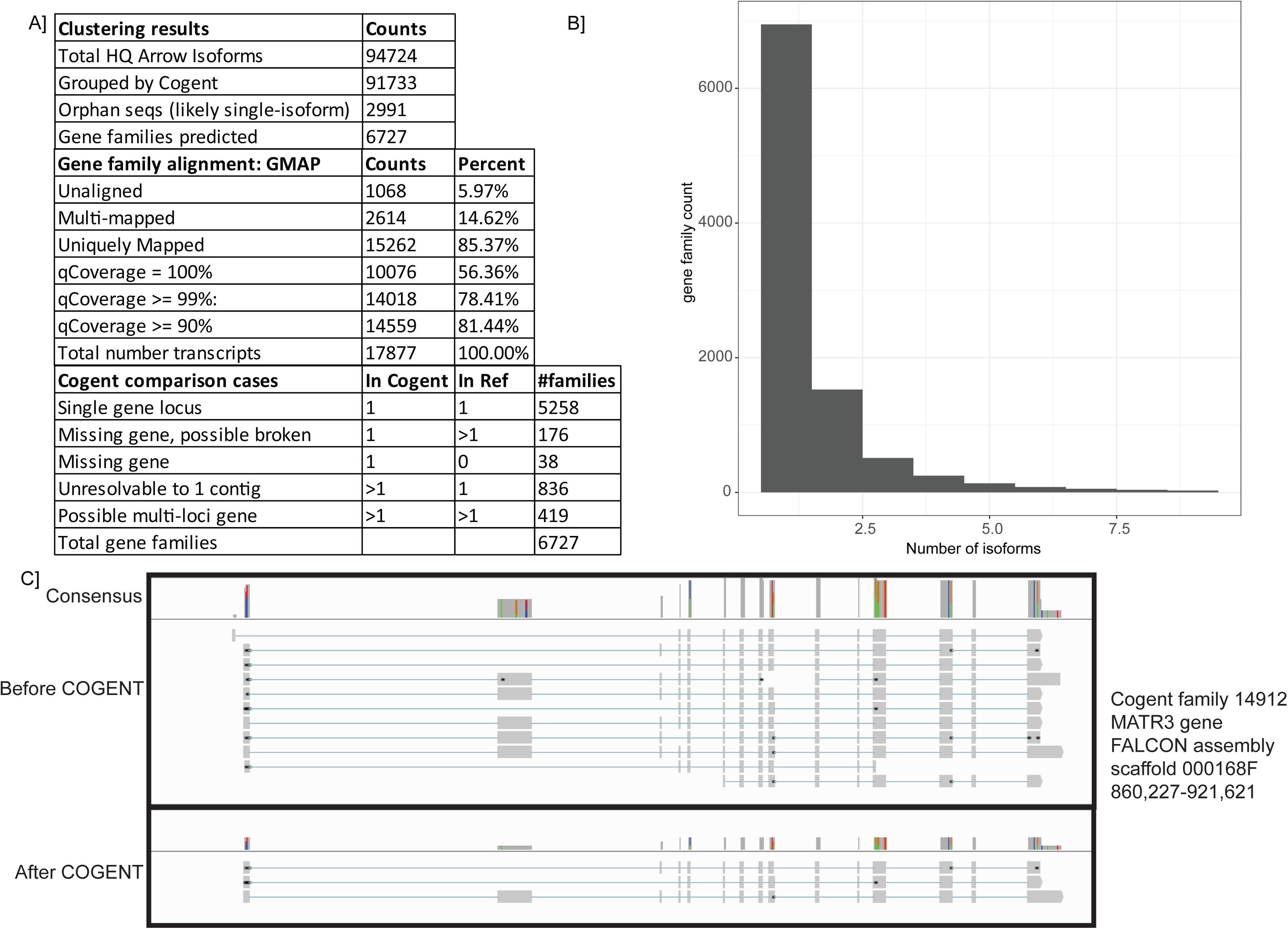
Reducing transcript redundancy and predicting gene families with COGENT. Cogent gene families predicted and classified by relationship to *Calypte anna* genome assembly **A**. Number of full-length reads which support each isoform reduced **B**. **C** IGV view of the MATR3 gene, which was reduced from 11 redundant reads to 3 unique isoforms using this pipeline

### Orthologous gene pair predictions and GO annotation show putative unique hummingbird orthologs

To examine protein sequence similarity and divergence between *Archilochus colubris* and other avian species, we used OrthoMCL, which generates reciprocal best hits from comparator species using BLAST all-vs-all, then clustering to group orthologous sequences for each pair of organisms (Li et al. 2003). OrthoMCL protein sequences were predicted using ANGEL, and 119,292 high quality sequences were put into this analysis. We compared our ruby-throated hummingbird, *Archilochus colubris,* to five other birds: *Calypte anna* (Anna's hummingbird) fellow member of the bee clade of hummingbirds, *Chaetura pelagica* (chimney swift) the closest available outgroup species to the hummingbird clade, and other bird species for which relatively well-annotated genomes and/or transcriptomes are available, *Gallus gallus* (chicken), *Taeniopygia guttata* (zebra finch), and *Melopsittacus undulatus* (budgerigar), as well as *Homo sapiens* (human), and *Alligator mississippiensis* (American alligator).

A matrix of ortholog pairings, with duplicate ortholog hits removed, shows counts of number of orthologous sequences for each species pair (Supplemental Table 3). Orthologs shared between ruby-throated hummingbird and a subset of the other species analyzed are illustrated in Figure 4A. Unsurprisingly, the largest amount of orthologs which pair closely to only one species, i.e., 1:1 orthologs, were found between Anna's and Ruby-throated hummingbird sequences. Surprisingly, the second-largest set was between chicken and ruby-throated hummingbird, as opposed to its closest outgroup species, *Chaetura pelagica.* This is likely due to the completeness of chicken transcriptome annotation, as chicken is the most well-studied avian species. Of the 596 unpaired *A. colubris* protein sequences, 190 paired most closely with *Calypte anna* when compared using BlastP and the majority of matches output (559/594) were less than 50 AA, only a fraction of the average sequence length.

**Figure 4.**
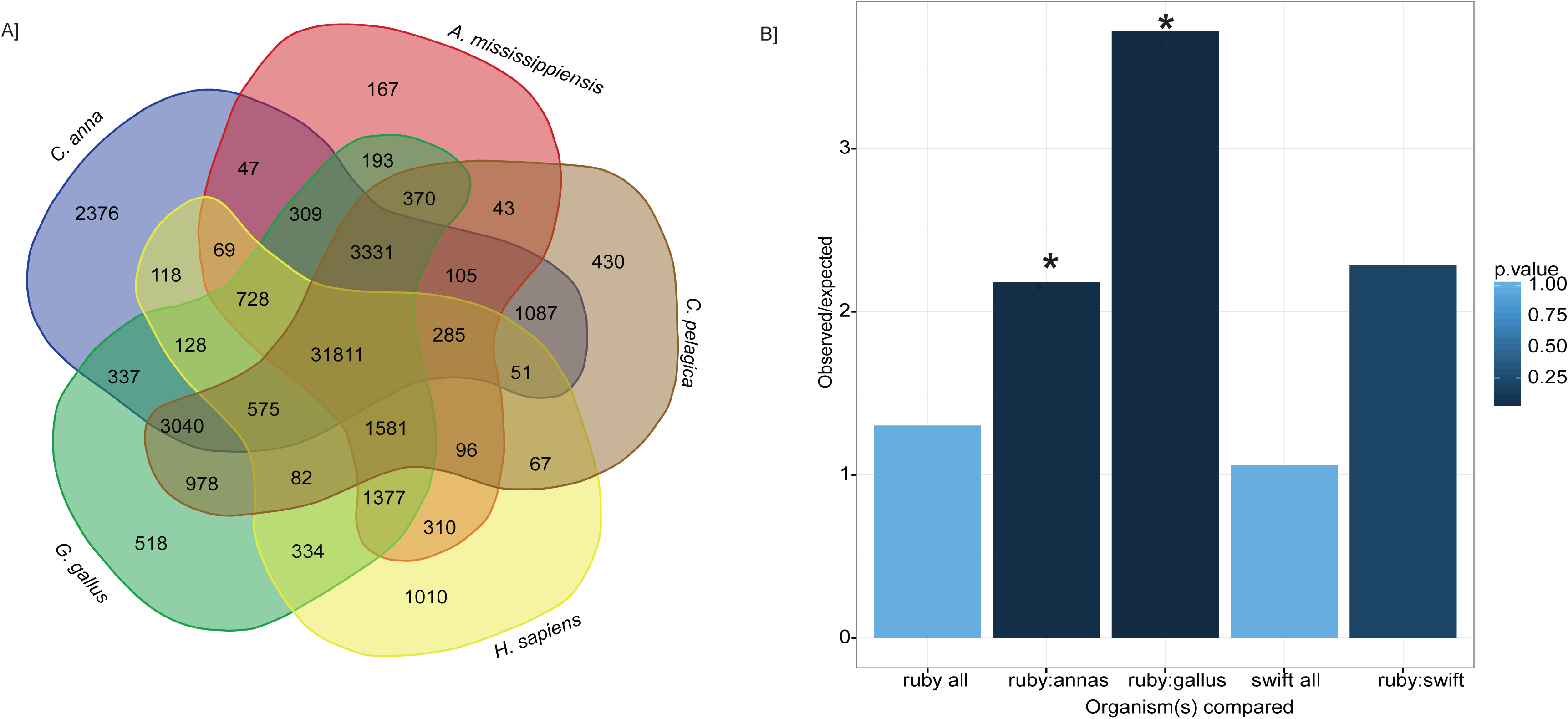
Orthology analysis. The transcriptomes of three birds (*Calypte anna, Gallus gallus, Chaetura pelagica*), one mammal (Homo sapiens) and one reptile (Alligator mississippiensis) were compared against *Archilochus colubris* using OrthoMCL to detect and compare similar sequences. **A** Venn diagram illustrating sequences with reciprocal blast hits between the given species and *A. colubris* is shown in A. Ortholog pairs unique to hummingbirds *Calypte anna* and *Archilochus colubris* were selected for gene orthology (GO) annotation analysis, which revealed enzymes of many biological functions (**B**).

In order to more closely examine the identity of 1:1 orthologs in related hummingbird species, gene ontology (GO) annotation was performed on a specific set of orthologs which were shared between *Calypte anna* and *Archilochus colubris,* but not by the other birds included in the OrthoMCL analysis. This set of 2,376 protein sequences was run using BlastP and GO analysis performed by Panther (Mi et al. 2013, 2017). Additional datasets used for GO comparison included 1:1 orthologs for *Gallus gallus* and *A. colubris* (518), and *A. colubris* and *Chaetura pelagica* (430), as well as whole transcriptome data from *C. pelagica* and COGENT-collapsed dataset from our transcriptome (Supplemental Table 4, Figure 4B).

As the initial impetus for our investigation centered on the exceptional metabolism and energetics of hummingbirds, we focused our investigation on orthologs tagged as part of the “metabolic process (GO:0008152)” grouping. Of the 1444 orthologs identified in *Archilochus colubris* as part of this process grouping, 236 (16.3%) were unique to hummingbirds. Within this top-level grouping, the largest number of genes group under “primary metabolic processes” (GO:0044238)”. Of the 1240 orthologs identified within this grouping, 204 (16.3%) are identified as uniquely shared by our hummingbird species. Six GO biological processes are defined under the “primary metabolic processes”. Of these processes, the process with the highest proportion of identified *A. colubris* orthologs hitting as unique to the two hummingbird species is “lipid metabolic processes” (GO:0006629; 33 of 114 orthologs, 28.9%), which is significantly enriched relative to the comparative orthology databases of both chicken and human. Because we considered it likely that an enrichment in lipid metabolic genes could be a result of our dataset being from liver tissue, we compared enrichment with that of the entire COGENT predicted gene set from the ruby-throated hummingbird transcriptome, and found no significant enrichment. This suggests a higher degree of divergence within this class of enzymes than would be predicted statistically.

The proportion of identified genes within a biological pathway classified as orthologs unique to hummingbirds should not be taken as direct evidence of greater selection on proteins within that pathway. Yet, if neutral sequence divergence is assumed to be randomly accrued throughout a species' genome, then greater divergence in enzymes making up “lipid metabolic processes” suggests that closer examination of these proteins for evidence of functional, or even adaptive, divergence is warranted. A phylogenetically-informed analysis of ortholog divergence among taxa is necessary to establish a selection signature, which will become possible in the future with the advance of the B10K project (Zhang et al. 2015) and larger numbers of avian species in GO databases.

### Hepatic lipogenesis case study demonstrates utility of long-read transcriptomics data in evaluating biology

Given the apparent sequence divergence among enzymes involved in “lipid metabolic processes” hinted at by orthology and ontology analyses, we elected to more closely examine sequence divergence in enzymes comprising the lipogenic pathway. Lipogenesis, the process by which fatty acids are produced from acetyl-CoA and subsequently triglycerides are synthesized, involves several key enzymes which we examined in closer detail. We predicted that this pathway (Figure 5A) would be divergent in hummingbirds given their extraordinary metabolic demands. Eight enzymes involved in this pathway were examined for *Archilochus colubris, Calypte anna, Gallus Gallus, Chaetura pelagica, Alligator mississippiensis* and *Homo sapiens* (accession numbers and details given in Supplemental Table 5). Pairwise protein alignment scores are given in Supplemental Table 6 as well as illustrated in a heatmap shown in Figure 5B, and alignments in Supplemental Data 1. Interestingly, enzymes with higher identity between examined organisms are involved in the fatty acid synthesis arc of metabolism, while triglyceride synthesis enzymes tend to be less conserved (Figure 5A). While poor pairwise protein alignment between *A. colubris* and all examined species, such as with DGAT2, is suggestive of misannotation or splice variation in our transcriptome, cases with variable alignment identities provide interesting targets for further investigation.

**Figure 5.**
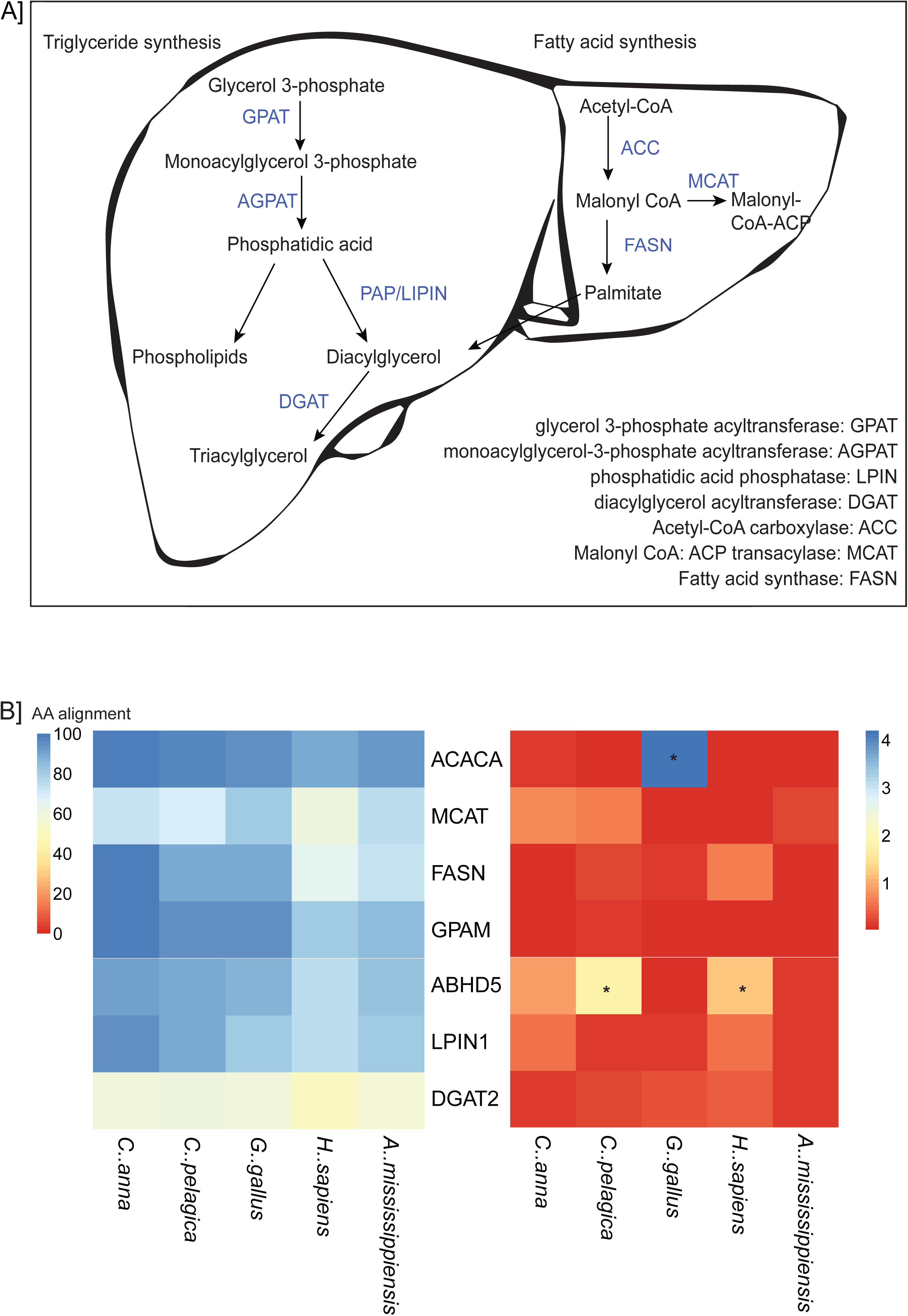
Pathway analysis of key enzymes in hepatic lipogenesis. **A** An overview of the relationship between the investigated genes and their roles in triacylglycerol, phospholipid and fatty acid synthesis, **B** a heat map illustrating percent identity of these proteins relative to *Archilochus colubris* predicted sequences, and nucleotide coding sequence conservation scores predicted using PAML (dN/dS), with genes dN/dS>1 starred.

In order to further investigate degree of conservation between key metabolic enzymes in hummingbirds and comparator organisms, we performed conservation analysis and determined ratio of nonsynonymous to synonymous codon changes (dN/dS) as a metric of positive selection, using pairwise alignments followed by codeml module in PAML4 (Yang 2007). These ratios are given in Supplemental Table 6. We found general conservation of these enzymes between organisms, with the exception of the 3’ and 5’ ends of alignments. These often had an extended or retracted coding sequence in the case of hummingbirds and *C. pelagica,* which could potentially be post-translational modification or selection on pathway regulation (Jacob and Unger 2007). Surprisingly, terminal sequence length was variable even between *C. anna* and *A. colubris,* which both belong to the closely-related Bee hummingbird taxid (McGuire et al. 2014). Variation in 5’ and 3’ length may also be an effect of the different methodologies used to produce these sequences, RNA sequencing for *A. colubris* and *G. gallus,* and ORF prediction from genomic data for the other organisms examined.

The averaged dN/dS values, while useful for comparison, are misleading when considered over the entire gene, as 3’ and 5’ variation can overshadow conserved motifs and vice versa, and pairwise comparisons are limited in scope. In addition, this type of analysis is ideal for very divergent sequences, and less informative for pairs of sequences that are highly similar (Kryazhimskiy and Plotkin 2008). Despite this, conservation analysis is still valuable and provides relative conservation metrics between these organisms for specific enzymes in a pathway of interest, as well as enabling identification of specific regions of high variation within coding sequences. Additionally, pairwise comparisons provide interesting observations, such as coding strand elongation in the 5’ region in *A. colubris* GPAM and GPAT4 (AGPAT6) (Supplemental Data 2). This information will be carried forth into future studies more closely examining enzyme structure, function and evolution.

Access to the transcriptome informs the investigation of biological processes and enables the formation of new hypotheses. This is exemplified by the serendipitous observation that hummingbird glucose transporter 2 (GLUT2) lacks a N-glycosylation site due to an asparagine to aspartic acid amino acid substitution. This missing glycosylation site was also seen in the available Anna's hummingbird genome. All class 1 glucose transporters studied in model vertebrates contain one N-glycosylation site located on the large extracellular loop of the protein (Joost and Thorens 2001). In GLUT2 the associated glycan interacts with the glycan-galectin lattice of the cell, stabilizing cell surface expression (Ohtsubo et al. 2013). Removal of the N-glycan of GLUT2 in rat pancreatic β cells results in the sequestering of cell-surface GLUT2 in lipid rafts and this sequestered GLUT2 exhibits a reduction in glucose transport activity by approximately 25% (Ohtsubo et al. 2013). This reduction in transport is thought to occur through interaction of the GLUT with lipid raft-bound stomatin (Ohtsubo et al. 2013; Zhang et al. 2001). In mammals, GLUT2 serves a glucose-sensing role in the pancreatic β cells and is required for the regulation of blood glucose through insulin and glucagon (Thorens and Mueckler 2010). The lack of N-glycosylation of GLUT2 may contribute to observed high blood glucose concentration in hummingbirds (Beuchat and Chong 1998).

## Conclusions

Our results have leveraged cutting-edge technology to provide a compelling first direct look at the transcriptome of this incredible organism. By using PacBio sequencing, we have been able to generate full length cDNA transcripts from the hummingbird liver. We subjected this data to a new and innovative pipeline, including steps to generate ORFs, merge transcripts to reconstruct gene families (COGENT), determine orthologs (OrthoMCL) and even measure evolutionary conservation and analyze genes.

Our data is remarkably representative of the transcriptome, capturing ~75% of metazoan universal orthologs, and ~50% of aves universal orthologs. This is an amazing diversity from a single tissue, given that many tissue specific genes are expressed at low levels if at all in liver. By subsetting our data, we show a saturation of the number of unique genes detected, suggesting that our data represents a nearly complete transcriptome of the hummingbird liver. Our data show a high degree of alignment to the existing *Calypte anna* dataset; even the reads which did not align with GMAP were subsequently mostly found to have a best match to *C. anna* with BLAST.

### Long read cDNA sequencing allows for assembly-free, less ambiguous isoform detection

Previous studies have shown that in a non-model organism, long read RNA sequencing improves detection of alternative splicing events nearly five-fold (Abdel-Ghany et al. 2016). Using full-length transcript data, we found alignment unnecessary to generate clear pictures of the gene isoforms and COGENT reconstructed gene contigs. The long reads negate the need for transcript assembly, a precarious analysis in the absence of a genome. The longest read identified was acetyl-coA carboxylase 1, with read length greater than 7kb, capable of spanning the length of the coding sequence. We have annotated our COGENT de novo transcriptome with the closest equivalent gene from NCBI nr/nt database (O'Leary et al. 2016) which, in combination with GO and OrthoMCL analyses, has improved transcriptome characterization. Although absolute functional assignments using GO annotation would not be ideal for this non-model species, comparing relative abundance of characterized orthologs between different datasets is useful for establishing a framework for forthcoming investigations.

### Transcriptome data provides insight into biological function of hummingbird liver

Often non-model organism protein sequences available in public repositories are translated from coding sequences predicted from whole genome data, not transcriptome data. While powerful, this approach for elucidating a transcriptome does not provide information regarding tissue-specific transcription, relative transcript abundance or isoform prevalence. In contrast, whole transcript mRNA sequencing from the liver tissue of *Archilochus colubris* has allowed us to clearly annotate gene families and the specific isoforms expressed in liver. In follow up studies, we will be able to compare the isoforms expressed between the liver and other tissues, e.g. pectoralis muscle.

### Orthology and gene ontology analysis give clues underlying uniqueness of extreme metabolism

Using our transcriptome, we have identified genes with unique orthologs in hummingbird as compared to other bird species, reptiles or even mammals. These genes showed a specific enrichment for pathways involved in lipid metabolism - suggesting that the hummingbird has evolved variants of these genes to achieve its high levels of metabolic efficiency.

### Polished transcriptome provides basis for future genomic and biological studies

Transcriptome data generated using the Iso-seq methodology, when coupled to sophisticated recently developed gene synthesis techniques (Kosuri and Church 2014), allows for simple generation of relevant isoforms for biochemical experiments. Some of the key metabolic enzymes identified from our work as being unique to either *A. colubris* or at most common to *C. anna* and *A. colubris* could be quickly cloned and expressed. Follow up studies will allow for biochemical studies of proteins generated directly from our transcriptome data, measuring their enzymatic properties, e.g. k_cat_ or V_max_, as compared to other avian or mammalian analogues (Suarez et al. 2009; Fernández M.J. et al. 2011; Suarez et al. 1988). Expressed proteins may also be used for structural biology studies, applying either x-ray crystallography or cryoEM to generate structural maps of the proteins, and examine how the structure compares to other analogues. Importantly, most of the glycolytic (hexokinase, phosphofructokinase) and lipogenic (acetyl-CoA carboxylase and pyruvate carboxylase) enzymes are highly evolutionarily conserved, and structural information exists for most (Lasso et al. 2014; Xiang and Tong 2008; Zhang et al. 2003; Aleshin et al. 1998; Kamata et al. 2004; Mulichak et al. 1998).

## Methods

### Sacrifice and RNA extraction

Wild adult male ruby-throated hummingbirds (*Archilochus colubris*) were captured at the University of Toronto Scarborough using modified box traps. Birds were housed in the University of Toronto Scarborough vivarium and fed NEKTON-Nectar-Plus (Nekton, Tarpon Springs, FL, USA) ad libitum. Birds were sacrificed after ad libitum feeding, and tissues were sampled immediately after euthanization using RNAse-free tools. A hepatopancreas tissue sample was collected from one bird. Tissue was homogenized at 4°C in 1 ml cold Tri Reagent using an RNase free glass tissue homogenizer and RNase free syringes of increasing needle gauge. We used 100 mg of tissue per 1 ml of Tri Reagent (Sigma-Aldrich, St. Louis, MO, USA), and chloroform extraction was performed twice to ensure quality. RNA was precipitated, centrifuged down, washed with ethanol, vacuum dried and eluted in RNAse free water. DNAse I digestion and spin column cleanup were performed. RNA concentration and RIN were determined with RNA Bioanalyzer (Agilent).

### Sample preparation and sequencing

Pacific Bioscience's Iso-Seq sequencing protocol was followed to generate sequencing libraries (Thomas et al. 2014). The Clontech SMARTER cDNA synthesis kit with Oligo-dT primers was used to generate first and second-strand cDNA from polyA mRNA. After a round of PCR amplification, the amplified cDNA was size selected into 4 size fractions (1-2kb, 2-3kb, 3-6kb, and 5-10kb) to prevent preferential small template sequencing, using the Blue Pippin (Sage Sciences). Additional PCR cycles were used post size-selection to generate adequate starting material, and then SMRTbell hairpin adapters were ligated onto size-selected templates. Each of the 4 size fractions was sequenced on 10 SMRT Cells, for a total of 40 SMRT Cells. Sequencing was performed by the JHU HiT Center using P6-C4 chemistry on the RSII sequencer.

## Analysis Methods

### Data processing, isoform clustering and sorting using the DNANexus Pipeline

The SMRTanalysis 3.1 software (https://github.com/PacificBiosciences/SMRT-Analysis) and IsoSeq pipeline were employed using a DNANexus interface. Raw sequence files produced from the Pacbio RSII (bax.h5, bas) were converted into BAM files using bax2bam, zmws_per_split 3. Circular consensus sequence (CCS) was generated from subread BAM files, parameters: min_length 300, max_drop_fraction 0.8, no_polish TRUE, min_zscore −9999, min_passes 1, min_predicted_accuracy 0.8, max_length 15000. CCS.BAM files were output, which were then classified into full length and non-full length reads using pbdassify.py, ignorepolyA false, minSeqLength 300. Non-full length and full-length fasta files produced were then fed into the cluster step, which does isoform-level clustering (ICE), followed by final Arrow polishing, hq_quiver_min_accuracy 0.99, bin_by_primer false, bin_size_kb 1, qv_trim_5p 100, qv_trim_3p 30.

### Aligning to reference using GMAP

We aligned with GMAP (Wu and Watanabe 2005) version 2016-09-23 with parameters -f samse -n 0 -z senseforce against Calypte anna genome (Zhang et al. 2014).

### Assessing transcriptome completion using BUSCO

BUSCO v2.0 BETA (https://gitlab.com/ezlab/busco, (Simão et al. 2015), accessed 9/8/2016) was used to benchmark transcriptome completion, by checking for essential single copy orthologs which should be present in a whole transcriptome dataset for any member of the given lineage. We used both Metazoan and Aves lineages (ortholog sets) to examine transcriptome completion.

### Gene family prediction and reducing transcript redundancy using COGENT

COGENT (coding genome reconstruction tool) v1.3 (https://github.com/Magdoll/Cogent) uses k-mer similarity profiles in order to partition full length coding sequences into gene families, after which it reconstructs contigs containing the full coding region.

### Orthologous gene prediction using OrthoMCL

To examine protein sequence similarity and divergence between *Archilochus colubris* and other avian species, we used OrthoMCL (http://genome.cshlp.org/content/13/9/2178.full). OrthoMCL works by performing an all-vs-all BlastP comparison of input sequences, and determining reciprocal best hit of all input pairings (cut off at e-5 and >50% match). Putative orthologs are the reciprocal best hits between species, while putative paralogs constitute reciprocal better hits within species. A normalized similarity matrix, followed by Markov clustering, produces ortholog groups. We compared our ruby-throated hummingbird *Archilochus colubris* to five other birds-*Calypte anna* (Anna's hummingbird, release date 2014-04-24, ftp://climb.genomics.cn/pub/10.5524/101001_102000/101004/Calypte_anna.pep.gz), *Gallus gallus* (chicken, Galgal5, ftp://ftp.ncbi.nih.gov/genomes/Gallus_gallus/protein/protein.fa.gz), *Chaetura pelagica* (chimney swift, release date 2014-04-24,ftp://ftp.ncbi.nih.gov/genomes/Chaetura_pelagica/protein/protein.fa.gz), *Taeniopygia guttata* (zebra finch, taeGut3.2.4, ftp://ftp.ensembl.org/pub/release-85/fasta/taeniopygia_guttata/pep/Taeniopygia_guttata.taeGut3.2.4.pep.all.fa.gz), and *Melopsittacus undulatus* (budgeriger, melUnd1, ftp://climb.genomics.cn/pub/10.5524/100001_101000/100059/BGIMUN1.120628.gene.withUTR.pep), as well as human (*Homo sapiens*, hg38, ftp://ftp.ensembl.org/pub/release-85/fasta/homo_sapiens/pep/Homo_sapiens.GRCh38.pep.all.fa.gz) and American alligator (*Alligator mississippiens,* allMis0.2/1, ftp://ftp.ncbi.nih.gov/genomes/ANigator_mississippiensis/protein/protein.fa.gz).

### GO analysis using PANTHER

Orthologs shared between *Calypte anna* and *Archilochus colubris,* but not with other birds included in OrthoMCL analysis, were examined more closely using gene ontology (GO) analysis. BlastP with default alignment settings was used to determine the top three putative hits for each ortholog. Genbank accession numbers were converted in a universal gene symbol using BioDB (https://biodbnet-abcc.ncifcrf.gov/db/db2db.php, (Mudunuri et al. 2009). Gene symbols were then run through Panther (http://pantherdb.org/, (Mi et al. 2017), producing GO terms for the input orthologs.

### Conservation analysis using PAML

PAML4.9c (Yang 2007) was used in order to estimate pairwise conservation between ruby-throated hummingbird and the five species (*Archilochus colubris, Calypte anna, Gallus Gallus, Chaetura pelagica*, *Alligator mississippiensis* and *Homo sapiens*) used as comparators in pathway analysis. Pairwise alignment was performed using CLUSTALX2.1 with PHYLIP output. Codeml module was used, runmode= −2, and default settings. dN/dS was estimated using Nei and Gojobori (Nei and Gojobori 1986). The mRNA sequences were then translated to protein using ExPasY, and these proteins were aligned using CLUSTAL, alignment scores were recorded.

### ORF prediction and protein translation using ANGEL

The ANGEL pipeline (https://github.com/PacificBiosciences/ANGEL), a long read implementation of ANGLE (Shimizu et al. 2006) was used in order to determine protein coding sequences from cDNAs. ANGEL consists of three primary stages: dumb ORF prediction, classifier training, and prediction. Dumb open reading frame (ORF) prediction, which produces all six possible open reading frames per given transcript, was run for all transcripts with a minimum length of 300 amino acids. Training involves the creation of a random subset of non-redundant transcripts, which is then used to create a classifier pickle file implemented in the prediction stage. Prediction then outputs the most likely ORF (minimum peptide length of 50 amino acids) based on length and coding potential of each given sequence. We performed this analysis on both our HQ polished isoform (HQD) and all sequences datasets (ASD).

This resulted in 119,292 HQD and 1,061,147 ASD peptide sequences, with a size distribution comparable to human, chicken, swift and alligator, with a mean amino acid length of roughly 500 AA, and a long tail.

## Data Accession

Filtered fastq of clustered CCS reads deposited in SRA accession number SRP099041. Predicted coding sequence and annotations, peptide and untranslated region data are available at Zenodo 10.5281/zenodo.311651. Genbank submission is in progress. All other data available upon request.

## Acknowledgments

Pacific Biosciences for reagents and SMRTcells as well as technical support. M. Schatz, E. Jarvis, J. Korlach, Y. Guo for discussion. HFSP grant #RGP0062/2016. Natural Sciences and Engineering Research Council of Canada Discovery Grant (#386466) to KCW.

## Disclosure Declaration

W.T. and R.W. have received travel funds to speak at symposia organized by Pacific Biosciences. Bulk of reagents for IsoSeq were provided by Pacific Biosciences.

